# Mutational scanning by multiplexed genome editing of the essential transcription termination factor Nrd1

**DOI:** 10.1101/2025.09.09.675026

**Authors:** Umberto Aiello, Kevin R. Roy, Lars M. Steinmetz

**Affiliations:** Stanford University School of Medicine, Department of Genetics, Palo Alto, CA, USA; Stanford Genome Technology Center, Stanford University, Palo Alto, CA, USA

**Keywords:** mutational scanning, genome editing, Nrd1, transcription termination, CTD-interacting domain

## Abstract

Proteins operate through a few critical residues, yet most proteins remain uncharacterized at the deep molecular resolution, particularly within essential genes, where functional dissection is obstructed by lethality.

Here, we establish a platform for mutational scanning of essential genes at their endogenous *locus*, combining a repressible complementation system with multiplexed CRISPR-based genome editing in budding yeast. Our approach provides a generalizable framework for dissecting essential protein function *in vivo*, expanding the capacity to map critical residues underlying essential cellular processes. We applied this strategy to *NRD1*, encoding an essential RNA Polymerase II (RNAPII) termination factor and performed a systematic alanine scanning with near-saturation coverage. We discovered novel and unexpected lethal mutations in the CTD-interacting domain (CID), thus revealing an unanticipated importance for this domain. Overall, our results demonstrate the power of our mutation scanning platform to map critical residues underlying essential cellular processes.

## INTRODUCTION

Although composed of many amino acids, a protein’s function relies on just a handful of key residues. Identification of these functional amino acids is paramount to our deep molecular understanding and to our ability to design drugs, engineer proteins, and treat genetic diseases. This is particularly challenging for essential proteins, whose limited amenability to manipulation is compounded by their requirement for cell viability.

Structural studies can sometimes offer clues when key residues cluster in solvent-exposed pockets, but they fall short when these are dispersed or buried in the protein fold. Nevertheless, such residues can still play critical roles for protein function, including mediating allosteric changes (Montserrat-Canals et al. 2025). On the other hand, unbiased functional studies offer a valuable direct readout of protein function, and are especially convenient when coupled to a readily measurable phenotype. Scanning mutagenesis is possibly the most high-throughput and unbiased approach, enabling the probing of every single residue in a protein of interest (Araya & Fowler 2011). Traditionally, past endeavors have often relied on ectopic expression systems, due to the challenges of precisely editing the endogenous genome. However, these systems do not take into account factors such as expression level, copy number, and genomic context, all of which can impinge on phenotypes.

We have previously developed MAGESTIC, a CRISPR-Cas9 based technology, that enables the editing of the budding yeast genome at scale (Roy et al. 2018, 2024). In *S. cerevisiae*, Double-Strand Breaks (DSBs) are repaired using Homology-directed Repair (HR) rather than Non-Homologous End Joining (NHEJ) (Jasin & Rothstein 2013). MAGESTIC leverages this natural feature by coupling single guide RNAs (sgRNAs), which direct Cas9 cleavage activity, to matching donor templates carrying the intended edits. To improve editing efficiency, we employ active donor recruitment, and we insert a random, unique barcode at a constant *locus*, to simplify the readout by pooled growth and amplicon sequencing. We reasoned that such methodology offers an unprecedented opportunity for performing mutational scanning of endogenous genes. Nevertheless, this strategy would suffer from challenges when targeting essential proteins, since key residues can solely be inferred by indirect observations, i.e., by the inability to recover viable cells when mutating a functionally relevant amino acid. Notably, it is impractical to exclude that such mutations are not efficiently generated, and in-depth molecular studies are often conducted with readouts that go beyond survival. To overcome such limitations, we coupled MAGESTIC to a synthetic repressible complementing system with the ability to maintain otherwise lethal mutations by controlled expression of a recoded target gene copy. Here, we provide a proof of concept for the application of such a framework in the context of an alanine scanning of the essential *NRD1* gene.

Nrd1 is a relatively well-known protein, with biochemical and *in vivo* evidence accompanied by a partially solved structure. Nrd1 is a 64 KDa RNA-binding protein with homology to the mammalian SCAF8 and SCAF4 anti-terminator factors (Yuryev et al. 1996). All these proteins hold an N-terminal CTD-interacting domain (CID), a single RNA recognition motif (RRM) and a segment enriched in alternating charged residues, containing several arginine-glutamate and arginine-serine (RE/RS) dipeptides (Figure 1A). Functionally, Nrd1 is a subunit of the Nrd1-Nab3-Sen1 (NNS) complex, which all together is responsible for the termination of RNA Polymerase II (RNAPII) transcription at Cryptic Unstable Transcripts (CUTs), snoRNA and snRNA units (Jensen et al. 2013). This defines a termination pathway distinct from the one acting at mRNA-encoding genes, which is instead based on the activity of the evolutionarily conserved Cleavage and Polyadenylation Factor-Cleavage Factor (CPF-CF) complex (Porrua & Libri 2015).

**Figure 1.**
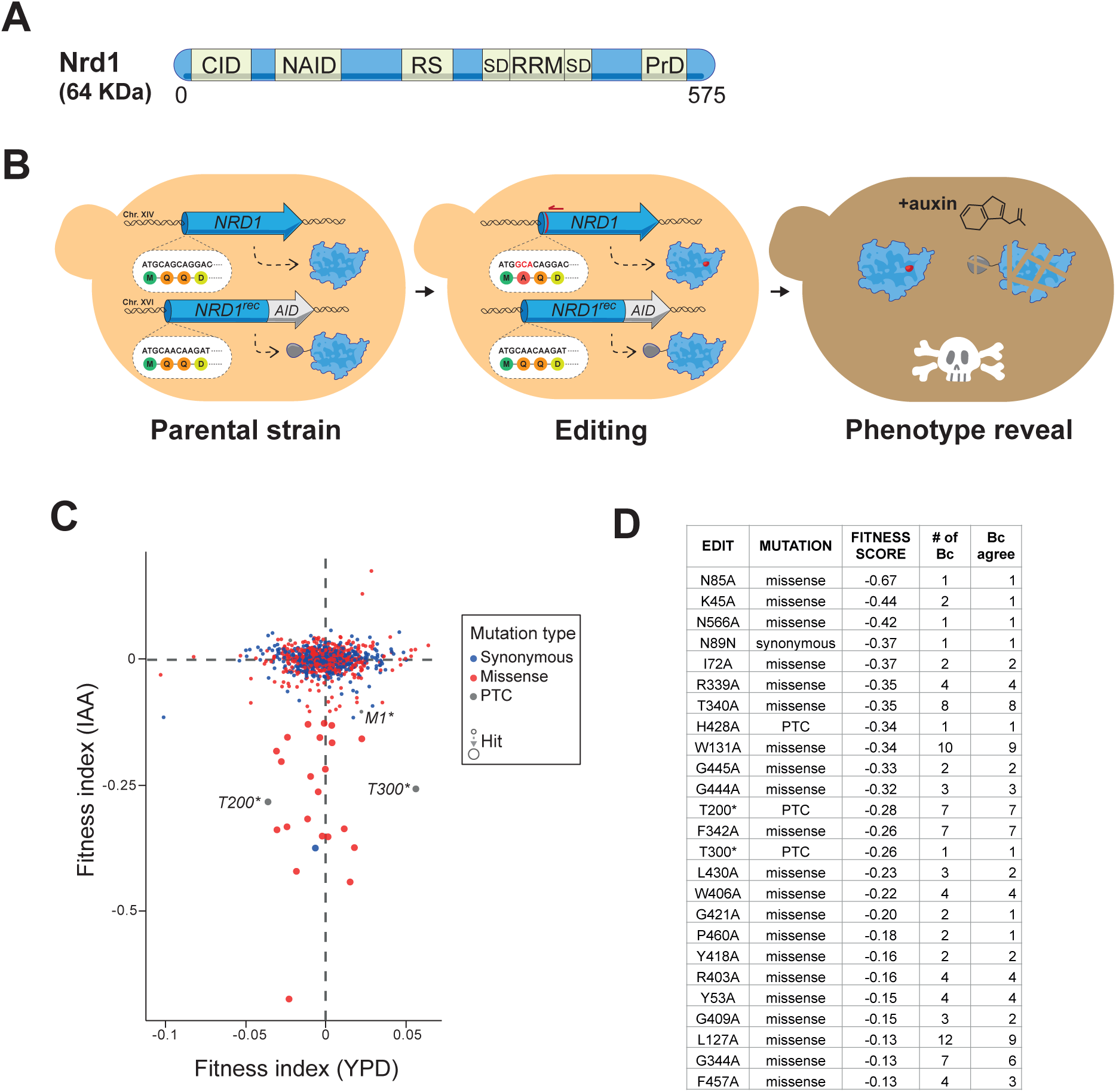
A platform to scan essential genes in yeast enables alanine scanning of the endogenous *NRD1* gene. **A.** Schematic representation of the functional domains along the Nrd1 protein. (CID: CTD-interacting domain; NAID: Nab3 interaction domain; RS: R/S rich region; SD: Split domain; RRM: RNA recognition motif; PrD: Prion-like domain) **B.** Overview of the platform to scan essential genes in yeast. First, a complementation system is introduced at a safe-harbor locus on Chr. XVI by recoding the sequence of the target gene. This second wild-type copy is expressed under the control of a degron system. During editing, CRISPR-Cas9 is guided only to the endogenous target gene, leaving the recoded wild-type copy to complement potential deleterious mutations. Finally, the phenotype associated with the introduced edit can be unveiled by auxin (IAA) mediated depletion of the recoded wild-type copy. **C.** Dispersion plot of the fitness scores of the edits computed over a 20-generation time course in the presence of IAA (phenotype exposed), and in YPD (phenotype complemented). **D.** List of the deleterious edits identified by filtering for the top 5% of effects, IAA-conditional effects and depletion by the end of the experiment. The number of barcodes (Bc) associated to each edit is indicated, as well as their agreement of the phenotype.

In the most widely accepted model of NNS termination, Nrd1 and Nab3 are recruited to the elongation complex (EC) by docking on the non-coding nascent RNA, via the recognition of specific motifs (Arigo et al. 2006; Thiebaut et al. 2006). The early binding of the Nrd1-Nab3 heterodimer was proposed to promote the recruitment, and the specificity, for the poorly abundant Sen1 helicase, via an interaction that involves Nrd1 CID and Sen1 Nrd1-interaction motif (NIM) domains (Han et al. 2020; Zhang et al. 2019). Sen1 subsequently evicts RNAPII from the DNA using its ATP-dependent translocase activity (Porrua & Libri 2013). Interestingly, disruption of the NIM region did not prevent the recruitment of Sen1 to the EC, nor lead to major termination defects at NNS targets (Han et al. 2017). Thus, the molecular function of Nrd1 remains elusive, yet the body of information provided by previous work makes it an ideal candidate for testing our strategy: known mutations are expected to be identified, while leaving a window open for discovering new biological insights.

## RESULTS

### A recoded complementation system enables editing of essential genes

To build our complementation system, we inserted a recoded copy of the target gene *NRD1* at a safe-harbor site located on the right arm of chromosome XVI (Figure 1B) (Flagfeldt et al. 2009). To recode, we preserved the protein coding potential, while altering the underlying DNA sequence as extensively as possible by taking advantage of the redundancy of the codon code. Thus, this second copy of the essential gene is shielded from the action of our CRISPR-based editing system and provides a supplementary wild-type protein to sustain survival even after insertion of otherwise lethal mutations into the target gene. Additionally, we appended an auxin-inducible degron tag to the recoded protein and introduced the Tir1 E3 ligase, required for the functionality of the degron system (Nishimura et al. 2009). Treatment with auxin or analogs (such as Indole-3-Acetic Acid, IAA) triggers the rapid degradation of the recoded protein, thereby allowing a direct fitness assessment of the mutation carried by the target gene.

We generated such a system for our target gene *NRD1* and validated expression of the recoded protein (Nrd1^rec)^ by Western blotting (Figures S1A and S1B) and confirmed that treatment with IAA led to a rapid depletion of Nrd1^rec^.

To test the complementing function of Nrd1^rec^, we edited the endogenous *NRD1* gene using CRISPR-Cas9-induced homology directed repair to introduce either of two known mutations: *nrd1-F378A* and *nrd1-V379G* (Bacikova et al. 2014; Conrad et al. 2000), both of which disrupt the RNA Recognition Motif (RRM) of Nrd1, which is required for its function. Whereas *nrd1-F378A* is inviable, *nrd1-V379G* is a temperature-sensitive allele, which causes lethality only upon growth at 30°C, while preserving viability at the 25°C permissive temperature.

We successfully recovered edited clones and verified the presence of the intended mutations by PCR and sequencing of the *NRD1* locus. The mutant cells did not display reduced fitness in standard growth conditions, demonstrating the complementing effects of Nrd1^rec^ (Figure S1C). However, treatment with IAA promptly induced a strong growth defect in cells carrying the *nrd1-F378A* mutation, whereas *nrd1-V379G* mutants, as expected, additionally required incubation at non-permissive temperature to exhibit lethality.

We concluded that our system efficiently complements lethal mutations in endogenous genes, and enables their phenotypic analysis on demand, even under conditional contexts.

### Comprehensive alanine scanning of *NRD1*

Since we did not intend to improve traits, and to maintain reasonable complexity in our library, we chose to forgo saturation mutagenesis and instead opted for an alanine scanning of the Nrd1 protein. Due to its chemical nature, alanine is unlikely to disrupt the folding of the protein, while only depleting the contribution of the native side chain (Cunningham & Wells 1989). For residues that are naturally alanine, we substituted glycine. We targeted each of the 573 positions (excluding the first and last codons) with two distinct alanine mutations, differing for RNA guide and donor, with this latter also containing several synonymous changes at the target site to prevent re-cleavage and targeting of the donor. To control for editing effects, we also designed a synonymous substitution for each missense mutation. Additionally, we included a small set of edits yielding Premature Termination Codons (PTCs) at different intervals to be used as positive controls for loss-of-function. In total, we designed 2309 edits. We employed the recently described MAGESTIC 3.0 approach (Roy et al. 2024) to introduce such edits in the yeast genome, with high efficiency and uniformity. After recovery of the edited cells, we assessed library composition. We detected edits that spanned 546 out of 573 available positions, thus covering ∼95.3% of the protein (Supplementary Figure 1D). When considering only missense mutations, we recovered edits at 525 positions (∼91.6% coverage). Of these, 478 positions (83.3% coverage) were represented by both a missense and a synonymous mutation. We observed >450 positions represented by more than 1 barcode, and >400 by more than 2 barcodes (Supplementary Figure 1E). As observed in Supplementary Figure 1F, we recovered a rather uniform distribution of edit along the gene, for both missense and synonymous mutations. Notably, only one region of 9 amino acids (532-540) failed to be edited. Since neither synonymous or missense edits were recovered in this region, it is unlikely that this depletion is caused by dominant-negative effects, and it is rather due to the high repetitiveness of this segment, which impacts editing efficiency (see Discussion).

Based on these observations, we conclude that our approach enables mutational scanning at endogenous *loci* and achieves near-saturation coverage.

### Fitness profiling identifies essential residues in Nrd1

To assess the impact of each mutation on the cellular fitness we tracked barcode abundance across a 20-generations time course, in triplicate, and performed in parallel in presence and absence of IAA. We computed a fitness index by fitting a regression line to the log_2_ fold change of the normalized barcode counts over time (see more in Materials and Methods) (Supplementary Figure 1G). We observed strong correlation scores among replicates (Supplementary Figure 1H).

We next called detrimental mutations by looking for edits that fell within the top 5% of effects, exhibited at least a three-fold greater impact in the presence of IAA compared to the control condition, and were depleted by the end of the time course. These criteria allowed us to identify a set of 25 positions (Figures 1C and D and Supplementary Table 1). Of these, 22 were associated with barcodes for missense mutations, 2 with PTCs, and 1 with a synonymous edit. 13 out of 17 of these hits were supported by multiple barcodes, with a remarkable agreement. The barcode count profile and regression analyses of these mutations are shown in Supplementary Figure 2.

To help visualize and interpret these detrimental mutations we generated heatmaps of the fitness score for each residue and displayed them along their position in the 2D dimension (Figure 2A). Lethal missense mutations were evident in the RRM region, which was expected since several inviable RRM mutants have been previously described and the whole domain is required for survival (Bacikova et al. 2014; Conrad et al. 2000). To our surprise, we also identified a second cluster of missense mutations that caused lethality, located in the CTD-interacting domain (CID). The latter is not required for viability (Steinmetz & Brow 1998) and its deletion only partially affects NNS termination, mostly by lowering its efficiency (Heo et al. 2013; Tudek et al. 2014). Together, these results confirm that our assay reliably detects the known essentiality of the RRM and, unexpectedly, uncover a second cluster of lethal substitutions in the CID region.

**Figure 2.**
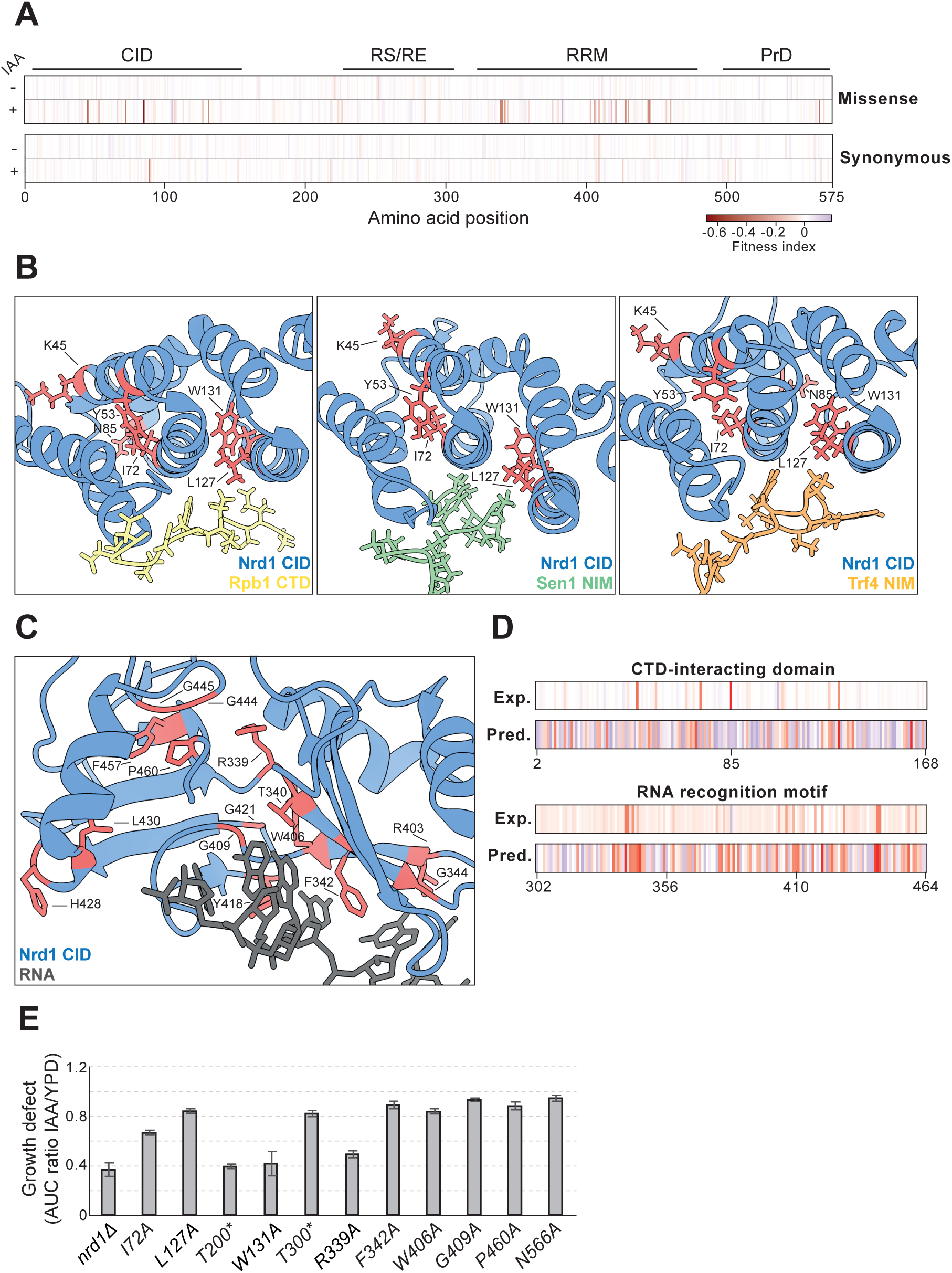
Alanine scanning of Nrd1 reveals detrimental mutations in the RRM and the CID domains. **A.** Heatmaps of the fitness score for the edits at the indicated position along Nrd1. Both missense and synonymous edits are shown, in presence and absence of IAA. The positions of the functional domains of Nrd1 are indicated on the top. B. Rendering of the structures of the CID domain bound to an Rpb1 (RNAPII) CTD repeat (left) (PDB: 2LO6), the Nrd1-interaction motif (NIM) of Sen1 (center) (PDB: 6GC3) and the NIM of Trf4 (right) (PDB: 2MOW). Lethal residues of Nrd1 identified in this study are highlighted in red. **C.** As in B, but for the RRM motif bound to an RNA (PDB: 5O20). **D.** Heatmap of the predicted (Pred.) instability of the CID (top) and the RRM (bottom) of Nrd1 using the FoldX AlaScan suite. The experimental fitness scores (Exp.) of these regions are also displayed. **E.** Growth defects for the indicated strains in the presence of IAA. The defect was computed as a ratio (IAA/YPD) of the area under the curve (AUC) of the normalized OD density over 6 generations of actively growing yeast cells.

### Structural mapping and validation of RRM and CID mutations

We next projected the residues identified as lethal onto available structures of the Nrd1 RRM (Franco-Echevarría et al. 2017) and CID bound to their relative partners (Kubicek et al. 2012; Tudek et al. 2014) (Figures 2B and C). For each structure, substitutions scored as detrimental in our screen were highlighted. In the RRM, these residues are predominantly located on the canonical RNA-binding face, whereas in the CID, they concentrate in the hydrophobic core of the folded domain, apart from L127, the only residue that makes contacts with the CTD, and Sen1 and Tfr4 NIMs. These projections indicate that lethal substitutions cluster in defined structural contexts.

We next compared our results with *in silico* alanine scanning, to determine if the observed effects could be explained by the predicted destabilization of these domains, using the FoldX suite (Schymkowitz et al. 2005) (Figure 2D). We obtained dissimilar results for the two domains: while the RRM displays a good level of agreement in between experimental fitness cost and predicted destabilization, the CID did not reveal a similar tendency, suggesting that additional mechanisms other than disruption of stability may come into play to explain the detrimental effect of the mutations in this domain. It is worth to note, that we also did not detect effects for many positions predicted to be highly detrimental in the CID, highlighting the limitations of inferring functional relevance solely from destabilization predictions.

The clustering of detrimental mutations in the RRM and a discrete region of the CID prompted us to examine representative substitutions in more detail through individual clone recovery and phenotypic assays. We performed Recombinase Directed Indexing (REDI) (Smith et al. 2017) (see more details in Materials and Methods), obtaining 1342 individual clones from the pooled library. Among these, we found several detrimental mutations identified by the screen, including *T200\** and *T300\** PTCs, *I72A* and *W131A* in the CID domain and *R339A* and *G409A* in the RRM region. All these strains were sequence-verified to ensure the presence of the intended edit. Additionally, we also constructed a *nrd1* deletion which we complemented using the Nrd1^rec^ system, to use as a reference phenotype for a complete loss of function.

To gain deeper insights on the effect of each of these mutations, as well as to validate their phenotypes, we performed non-competitive growth assays in liquid culture (Figure 2E and Supplementary Figure 3). In these tests, CID mutants (particularly *I72A* and *W131A*), displayed a remarkable fitness decrease in the presence of IAA, comparable to the strongest RRM mutation, *R339A*, and a full loss-of-function. Additionally, the analysis showed that *T200\** and *T300\** PTCs exhibited diverging growth defects, with the latter associated with a much weaker detrimental effect, suggesting a partial function of the protein truncated at position 300.

### Analysis of expression and localization of *nrd1* mutants

The lethal outcome of detrimental mutations can arise from multiple mechanisms, broadly categorized as direct (loss of enzymatic activity, toxic gain-of-function, and disruption of critical interactions) and indirect (protein mislocalization, misfolding, partial instability, as well as impacts on gene expression or RNA stability). To identify mutations where lethality stems from indirect mechanisms, we inserted a GFP epitope at the N-terminus of the *nrd1* edited *locus* in a selection of lethal substitutions.

First, to determine if some of these mutants produce unstable proteins, we monitored their expression levels by Western blotting analysis. None of the mutants displayed differences in the expression levels, beside *R339A* and the PTCs, for which a decrease in expression was observed (Figure 3A). Moreover, the analysis shows that *T200\** and *T300\** substitutions are *bona fide* PTCs, with their protein size consisted with the expected truncations. We concluded that the surveyed mutations are unlikely to destabilize the protein, except for *R339A*.

**Figure 3.**
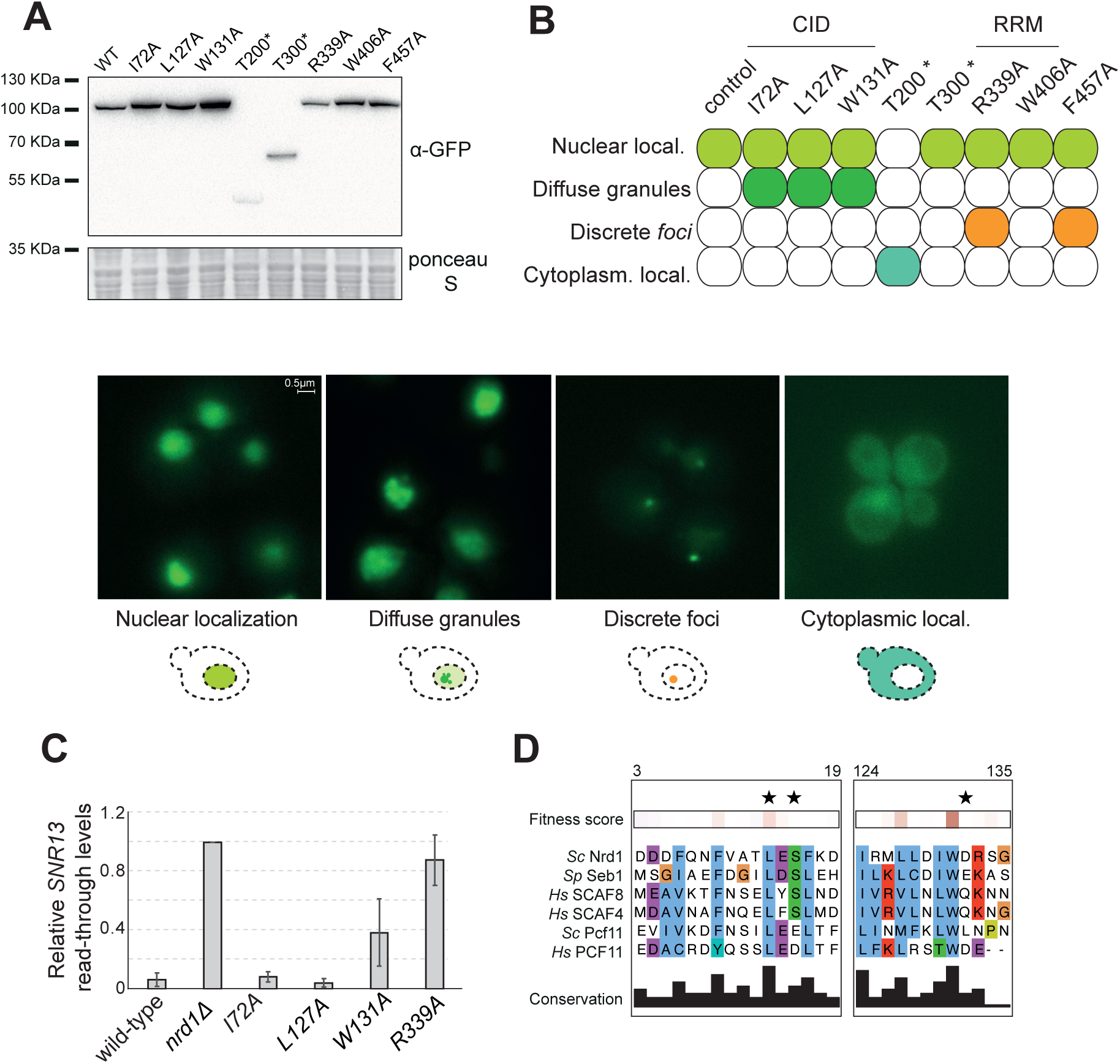
CID and RRM mutations produce lethality through distinct mechanisms. **A.** Western blotting analysis of GFP-tagged Nrd1 analogs. **B.** (Top) Summary table of the different phenotypes displayed by each strain; (bottom) representative microscopy fields and schematic representation of the different categories of Nrd1 localization identified. **C.** RNA quantification by RT-qPCR of the extended *SNR13* transcript in the indicated strains upon IAA treatment. **D.** Fitness score and alignment of the protein sequences of *S. cerevisiae* Nrd1, *S. pombe* Seb1, *H. sapiens* SCAF4 and SCAF8, *S. cerevisiae* Pcf11 and *H. sapiens* PCF11, in the indicated regions. Stars indicate positions in SCAF4 the mutation of which has been linked to neurodevelopmental disorders.

Finally, to verify the correct localization of the mutant proteins, we carried out microscopy analysis, taking advantage of the same GFP-tagged strains. We observed varied localization of the Nrd1 protein among the different mutants, yet striking similarities when mutations affected the same domain. We identified distinct phenotypes that we classified into four categories: diffuse nuclear localization, nuclear localization with diffuse granules, bright distinct *foci*, and cytoplasmic localization (Figure 3B and Supplementary Figure 4). The wild-type Nrd1 protein displays a strong and uniform nuclear signal. Instead, CID mutants, and particularly those with the strongest lethality, had a nuclear signal with a characteristic granularity. RRM mutants exhibited discrete, sharp and bright *foci*. Finally, the two PTCs had different localizations, with *T300\** being nuclear, and *T200\** completely cytoplasmic, thus likely explaining the dramatic difference in fitness cost of these two truncations. This also suggests that the region in between position 200 and 300 can mediate nuclear localization. We also conclude that CID and RRM mutants exhibit different localization patterns that might underscore distinct mechanisms causing their associated lethality.

### CID and RRM mutations produce lethality through distinct mechanisms

It is widely agreed that the lethal effects caused by malfunction of any NNS subunit are due to sudden deficiency of RNAPII transcription termination (Arigo et al. 2006; Schulz et al. 2013; Thiebaut et al. 2006). This leaves elongation complexes unchecked, and unfolds in dramatic downstream alterations on the transcriptome (Aiello et al. 2022; Schaughency et al. 2014). To investigate whether the *nrd1* mutations we identified in our screen followed the same logic, we sought transcription termination defects by inspecting RNAPII read-through at a well characterized NNS target, the *SNR13* gene. Here, lack of NNS termination results in an extended and partially stable RNA molecule that can be readily detected by reverse transcription and quantitative PCR (RT-qPCR). As shown in Figure 3C, the full depletion of Nrd1 recapitulates the expected major termination defects at *SNR13*. In a similar fashion, RRM mutations displayed a marked increase in read-through product. Conversely, the CID mutants surprisingly showed none or only minor termination defects. These results suggest that while RRM mutants fail to bind RNA and cause RNAPII read-through, the underlying lethality observed for CID mutants is not or only partially explained by termination defects.

## DISCUSSION

When it comes to their inner workings, proteins are intricate machines. Most residues support structural integrity, whereas only a few are directly involved in catalysis, binding or allosteric regulation. Not always these functional residues are obvious from sequence, structure or conservation alone, making their systematic identification critical for a deep molecular understanding. Here, we present a platform for the mapping of amino acid positions that are crucial for the protein’s function through mutational scanning at endogenous *loci* in budding yeast. Our setup combines multiplexed genome editing with a complementation system, enabling the generation of large libraries of analogs, as well as to mutate essential proteins without the confounding preventive loss due to detrimental substitutions. The utility and relevance of our approach is here demonstrated by our findings for the *NRD1* gene. We revealed that its CTD-interacting domain (CID), previously considered a non-essential binding surface, plays a central role in mediating Nrd1 function.

### On mutational scanning via genome editing

Scanning mutagenesis holds the potential to be one of the most powerful approaches to undertake the challenge of identifying critical residues in a protein. While technical difficulties have historically limited its application, advances in DNA synthesis, sequencing and library design have partially relieved some constraints. The effectiveness of this approach is clearly demonstrated by studies where mutational scanning has provided insights that would have been difficult to obtain otherwise, for example the functional annotation of 4000 SNVs in BRCA1 (Findlay et al. 2018), the identification of antibody-escape mutations in the SARS-CoV-2 receptor-binding domain (Starr et al. 2020), and the discovery of functional hotspots in G-protein–coupled receptors that were not evident from structural data alone (Howard et al. 2025). Nonetheless, a major effort is still needed to seamlessly perform mutational scanning at endogenous *loci*. Until very recently, virtually the totality of previous efforts in mutational scanning or related approaches have been carried out using ectopic expression systems. Yet, factors such as dosage, copy numbers and even genomic context matter, and they impinge on the final readout (Cooper et al. 2024; Inoue et al. 2017), as observed for certain endogenous TP53 variants that displays opposite phenotypes compared to exogenous overexpression systems (Gould et al. 2023). However, editing an endogenous gene can only be accomplished with caveats, such as by combining a cell context that tolerates HDR, donor design, and functional selection pressure. To overcome these limitations, we combined the use of MAGESTIC with a complementation system, and delivered a universal platform that can be used to scan any protein, regardless of their essentiality, and without further adjustments (Figure 1). By definition, dominant negative mutations cannot be complemented by our system. However, editing of the recoded copy upon constant IAA treatment should enable the safe introduction of such mutations, whose effects can then be directly visualized by removal of IAA. Thus, by simply switching the editing target (i.e., endogenous vs recoded copy) both recessive and dominant negative mutations can be studied using the same platform.

We demonstrate the feasibility of our strategy using Nrd1 as a test case. Nrd1 is approximately the size of an average yeast protein, and therefore, is a good example of the task of performing scanning mutagenesis in this organism. We demonstrate to reach near-saturation mutagenesis, with ∼95% of the protein edited. It is worth to note that a 9-residue stretch (positions 532-540) was entirely absent in our library for both missense and synonymous mutations, despite being represented prior to editing (Supplementary Figure 1J). As this deficit is not linked to mutation identity, we infer decreased editing efficiency at this specific segment. Its underlying coding sequence is highly repetitive, encoding a P/Q-rich stretch that constitutes Nrd1’s prion-like domain. In a recent survey of the genome-wide editing efficiency using MAGESTIC, we have demonstrated that repeated regions are prone to cause structural variants (SVs) upon CRISPR-Cas9 cleavage, which can cause severe lethality (Li et al., under revision; https://apps.embl.de/score). In the future it will be interesting to test if “preventive” measures can alleviate such limitation of the editing system, for example by decreasing the complexity of the target sequence prior to editing.

### On the detrimental mutations identified in *NRD1*’s RNA recognition motif

By applying our framework to *NRD1*, which encodes an essential transcription termination factor in the Nrd1-Nab3-Sen1 (NNS) pathway (Jensen et al. 2013; Porrua & Libri 2015), we identified expected and unanticipated classes of mutations. In particular, alanine substitutions in the RNA recognition motif (RRM) were broadly detrimental, consistent with genetic and biochemical evidence that the RRM is required for viability and for recognition of ncRNA motifs at NNS targets (Bacikova et al. 2014; Conrad et al. 2000). A highly detrimental cluster of mutations was concentrated within the groove contacting the RNA (Figure 2C). This served as an internal validation, showing that our workflow recapitulates established structure-function relationships.

We observed a strong agreement with *in silico* modeling using the FoldX algorithm, with the most detrimental mutations from our screen overlapping positions that are predicted to lead to the strongest destabilization of the domain (Figure 2D). This suggests that the main determinant of the RRM function is the ability to fold in a stable conformation that can properly interact with the RNA. This is further support by our microscopy analyses, which showed dramatic alterations in the localization of Nrd1 in RRM mutants (Figure 2B). Such effects are exemplified by the *R339A* mutation, which is also predicted to cause strong destabilization of the RRM and that results in the formation of a bright nuclear *focus*, suggestive of aggregation. Notably, structural analyses of the RRM domain showed that R339 lies at the center of an intricate network of hydrogen bonds and salt bridges (Franco-Echevarría et al. 2017), thus explaining the key role of this position in mediating the proper folding of the domain. It remains unclear why protein aggregation does not take place in the cytoplasm, even before Nrd1 is imported in the nucleus. One possibility is that perhaps binding to RNA counteracts an intrinsic nature of the protein to phase-separate and aggregate, possibly driven by its prion-like domain. This hypothesis is supported by the fact that RNA-binding proteins with prion-like behavior have been shown to display aggregation levels that can be buffered by binding to RNA (Langdon et al. 2018; Maharana et al. 2018). Moreover, Nrd1 and Nab3 were shown to form nuclear speckles in nutrient depletion (Darby et al. 2012), and Nrd1 was shown to form amyloid-like filaments, providing a biochemical basis for self-assembly (O’Rourke et al. 2015).

### On the detrimental mutations identified in *NRD1*’s CTD-interacting domain

In this study, we made the unexpected discovery of lethal mutations in Nrd1’s CTD interacting domain (CID). It was previously shown that deletion of this domain did not cause lethality (Steinmetz & Brow 1998) and only diminished the efficiency of NNS termination (Tudek et al. 2014; Vasiljeva et al. 2008). Yet, we found mutations with a strong detrimental effect, comparable to the strongest mutation in the RRM we have identified (compare *W131A* or *I72A* to *R339A* in Figure 2E).

By examining transcription termination efficiency at a widely used NNS target we made an additional surprising finding. Indeed, CID mutants did not show the expected termination defects typically observed in NNS mutants (Figure 3C). While *W131A* displayed some termination defects, these are still 0.5-fold lower than what we observed for *R339A*, despite these two mutations being similarly deleterious. Other CID mutants, such as *I72A* and *L127A,* did not show any termination defect at *SNR13*. This suggests that the mutations in the CID compromise a specific function of Nrd1, which does not lead to disruption of NNS termination.

Our findings seem in sharp contrast with the previously described non-essentiality of the CID. Nevertheless, it is possible that the point mutations, by disrupting the hydrophobic core of the CID may lead to a partially misfolded protein, whereas the deletion, by removing the entire domain, leaves a “clear” truncated protein. However, our microscopy and Western blot analyses (Figure 3) did not hint to misfolding or aggregation for any of these proteins, including *W131A*, which is predicted to be a strong destabilizing mutation. Notably, and in contrast with what we had observed for the RRM, we did not observe a neat agreement between FoldX prediction and *in vivo* fitness cost (Figure 2D). Many positions that are predicted to cause strong destabilization are not found as detrimental in our screen, and conversely not all mutations that we identified as lethal are predicted to be destabilizing. Thus, lethality of CID mutations may not stem primarily from domain destabilization.

An alternative explanation to the lethality of the CID mutants is that they may lead to the formation of aberrant poised complexes, whereas this is not case for the deletion. In fact, it seems unlikely that the function that it is disrupted in these mutants is the mere interaction between RNAPII CTD and Nrd1, because we did not find detrimental effects for mutations that disrupt the CID binding interface, and this would be in agreement with the deletion of the domain not being lethal. The mapping of these lethal mutations in the 3D structure shows that they are instead typically located in the hydrophobic core of the domain (Figure 2B). *L127* is the sole position that directly contacts RNAPII CTD, but its mutation towards alanine is not predicted by FoldX to significantly change the binding energy (ΔΔG = 0.11).

NNS termination was suggested to be temporally coordinated through a molecular mimicry that is also based on the CID and its interaction with different partners (i.e., the CTD, Sen1, Trf4) (Han et al. 2020). We speculate that the identified lethal CID mutations may perhaps disrupt this coordination and, by an as-yet unclear mechanism, trap elongation complexes in a poised, nonproductive state. This hypothesis could explain the severe lethality despite the absence of overt termination defects. The peculiar localization phenotype we observed for these mutants might indeed reflect unphysiological chromatin processes. Interestingly, CID mutations in SCAF4, a protein that shares large homology with Nrd1, have been recently shown to lead to misfunction, and to underly a human pathological condition (Schmid et al. 2025) (Figure 3D), thus perhaps paralleling a mechanism from yeast to humans.

## MATERIALS AND METHODS

### EXPERIMENTAL MODEL AND SUBJECT DETAILS

#### Base strain description

*Saccharomyces cerevisiae* is the model organism used in this study. All strains used in this study derive from DHY214 developed at the Stanford Genome Technology Center. DHY214 is a BY4742-derivative and contains modifications that improve growth, respiration, and sporulation by exchanging the minor allele present in BY/S288C-based strains to the major allele present in all other *S. cerevisiae* strains (Harvey et al.). The full genotype of *DHY214* is *MATα his3Δ1 leu2Δ0 ura3Δ0 lys2Δ0 CAT5(91M) SAL1 MIP1(661T) HAP1 MKT1(30G) RME1(INS-308A) TAO3(1493Q)*.

#### MAGESTIC 3.0 background strain overview

The MAGESTIC 3.0 base strain yKR965 was previously described (Roy et al. 2024). Briefly, yKR965 was derived from DHY214 by standard CRISPR-Cas9 genome editing approaches with SpCas9 and guide RNA plasmids and linear donor DNA to introduce genomic modifications (DiCarlo et al. 2013). First, several markers were deleted to enable selection and counter-selection (*fcy1Δ his3Δ* [full CDS and terminator deletion] *can1Δ lyp1Δ arg4Δ*). Second, the barcode landing pad was inserted at *YBR209W* (yT177 modification; *natNT2*- *LYS2*-*CAN1*-lox66-SceI_site-*LEU2*-SceI_site-*kanR::ybr209wΔ*). This landing pad contains homologies for integration of the barcode present on the guide-donor plasmids (Roy et al. 2024). It also contains a loxP variant site (lox66) that allows for recombinase-directed indexing (REDI) (Smith et al. 2017). Finally, the guide-donor plasmid removal system was introduced. It is dual galactose- and anhydrotetracycline (aTc)-inducible SaCas9 system designed for removal of guide-donor plasmids from cells post-editing. The SaCas9 guide targets a synthetic site (denoted site “X”) present on the guide-donor plasmid backbone and is under the control of an *RPR1* promoter with *tetO* sites for modulation via the aTc-inducible WTC846 system (Azizoglu et al. 2021; Roy et al. 2024). SaCas9 and the guide were inserted at YNRCΔ9 (referred to as “site 16” in Flagfedt et al., 2009; yT170 modification; p*GalL*_SaCas9-p*RPR1*_*tetO*_HHR_SaCas9_X1_sgRNA::*YNRCΔ9*) and the WTC846 system was inserted at YORWΔ17 (referred to as “site 20” in Flagfedt et al., 2009; yT138 modification; p*RNR2*-*tetR*-*TUP1*-p*TDH3*_*tetO*-*tetR::YORWΔ17*). yKR965 also contains pS1381, a centromeric plasmid with *URA3, hphMX*, SpCas9 under constitutive *TEF1* promoter, the Ec86 retron reverse-transcriptase under the *ADH1* promoter, and the donor recruitment fusion protein MCP-LexA-FHA under the *DUT1* promoter (Roy et al. 2024).

### METHOD DETAILS

#### Generation of the complementation system

The complementation system was introduced into the MAGESTIC base strain yKR965 by assembling the following fragments via transformation-associated recombination upon CRISPR-Cas9 cleavage at *YPRCΔ15* on Chromosome XVI (referred to as “site 20” in Flagfeldt et al., 2009) :(i) a cassette encoding OsTIR1::3xMyc under the control of a constitutive *ADH1* promoter, amplified by PCR from pOsTIR1w/oGFP (Papagiannakis et al. 2017); (ii) a constitutively active ribosomal promoter (p*RPLB8*), amplified from wild type *S. cerevisiae* genome; (iii) a recoded *NRD1* gene, which sequence follows the logic described below and synthesized by Twist Bioscience; an AID::6xFLAG tag, amplified by PCR from pHyg-AID*-6FLAG (Morawska & Ulrich 2013). Each of these fragments carried homology regions to allow integration (first and last fragment) and assembly among all parts (all fragments).

Recoding was performed by replacing each codon with the other most frequent alternative codon, if any available, and never exchanging for rare codons.

After selection on CSM lacking uracil and histidine, individual clones were passaged to fresh plates and single colonies reisolated. Correct editing was verified by PCR and sequencing of the integration site by Oxford Nanopore Technology (Plasmidsaurus). The functionality of the system was tested in the presence and absence of 500 μM IAA by Western blot with an antibody to the FLAG tag and by growth inhibition of a strain containing a deletion of the endogenous *NRD1* coding sequence.

#### Integration of the editing machinery at the barcode locus

To enable simultaneous removal of the editing machinery and integration of the barcode from the guide-donor plasmid post-editing, we modified YLS30 by replacing *LEU2* at the barcode locus with SpCas9 from pS1381 and the Ec86 retron RT and a modified version of the donor recruitment machinery from pS1505 (yT196 modification; p*TEF1*-SpCas9-*hphMX*-p*ADH1*-Ec86 RT-p*DUT1*-MCP-EcKlenow DNA polymerase-LexA-FHA-*KlURA3*). The strain (designated yKR1026) was confirmed by PCR and sequencing of the integration site by Oxford Nanopore Technology (Plasmidsaurus).

### Multiplexed genome editing via MAGESTIC

#### Guide-donor library design

Guide-donor sequences were designed to target each codon in the *NRD1* coding sequence with either an alanine (or glycine in the case of alanine) or a synonymous substitution. For each alanine/glycine or synonymous substitution, two guides with a 3’ NGG protospacer-adjacent motif (PAM) for SpCas9 were selected. Additional synonymous substitutions adjacent to the target codon were introduced as necessary to prevent SpCas9 (re-)cutting of the edit and donor DNA. Scripts and input files for the library design are available at https://github.com/k-roy/MAGESTIC/tree/master/NRD1. As an example, an oligo designed to introduce an alanine at position 4 (*NRD1* D4A) is 5’-ctgcgattggcaggcgcgcc<u>TAAACATCCCATAATGCAGC</u>gtttgaagagc<u>CAAGCACGAGTTACAGGA AAGGAACCGGAAAGCAACAAACATACTAAACATCCCATAATGcaacaagcaGACGATTTTCA AAATTTTGTAGCTACCTTGGAATCATTCAAAGATTTGAAATCTGGTATT</u>cgatagacgactggaca gca-3’. The guide (20mer) and donor (129mer) sequences are underlined. They are separated by a BspQI cloning site (gtttgaagagc) and flanked by priming sites for library amplification, with subpool specificity provided by the reverse priming site (cgatagacgactggacagca). In the middle of the donor sequence, the edited codons are in lower-case. In this case, the target codon position for D is mutated to gca and contains two upstream synonymous substitutions for Q (caacaa). The *NRD1* guide-donor library consisted of 2308 sequences and was ordered as a single oligo pool at the 200mer length and 180k scale from Twist Biosciences.

#### Step 1 plasmid library cloning

The dried oligo pool was resuspended in 1X buffer TE (10 mM Tris buffer, pH 8.0 with 1 mM EDTA) to 20 ng/μL and kept at -80°C in 10 μL aliquots. The pool was amplified with KAPA HotStart Polymerase (Roche) in a 50 μL reaction with 1 μL of oligo pool (20 ng/μL; actual subpool concentration is ∼0.26 ng/μL due to the *NRD1* subpool occupying only a fraction of the oligo pool), 1.5 μL forward and reverse primers, and 25 μL of 2X KAPA HotStart master mix with the following program: 95°C 3 min, 17 cycles of [98°C 20 sec, 60°C 15 sec, 72°C 30 sec], 72°C 2 min. The forward primer is

5’-cgttcgaaacttctccgcagtgaaagataaatgatcggagctgcgattggcaggcgcgcc-3’

and the reverse primer is:

5’-CATTGAATCTGAGTTACTGTCTGTTTTCCTNNNNNNNNNNTGCTGTCCAGTCGTCTATCG-3’.

The N10 sequence appends each guide-donor oligo with a random barcode that is utilized for associating each donor with its observed guide after cloning. The reactions were cleaned by Ampure XP beads in a 1.8:1 bead solution:sample ratio and eluted in 25 μL buffer EB (Qiagen) with a concentration of 13.5 ng/μL as measured with the Qubit dsDNA High Sensitivity kit (Invitrogen). In parallel, the MAGESTIC 3.0 guide-donor plasmid pS1439 was cut with AscI and BspQI (NEB) and purified by gel extraction, yielding a concentration of 93.4 ng/μL by Qubit. 2 μL of cut pS1439 backbone and 8 μL of the amplified *NRD1* library were assembled with 10 μL of NEBuilder® HiFi DNA Assembly Master Mix in 20 μL total volume at 50°C for 1 hour and then placed on ice. 2 μL 3M NaOAc (pH 5.2) and 1 μL GlycoBlue™ Coprecipitant (15 mg/mL, Invitrogen) were added to the 20 μL assembly on ice, mixed well by pipetting, and combined with 160 μL 100% ethanol for a standard ethanol precipitation. Pellets were resuspended in 3 μL H2O and 1 μL electroporated into 20 μL E. cloni 10G Elite Electrocompetent Cells (LGC Biosearch™ Technologies) with standard parameters as specified by the manufacturer (cuvette with 1.0 mm, 10 μF, 600 ohms, 1800 volts). Cells were resuspended in 1 mL 37°C pre-warmed E. cloni Recovery Medium and recovered for 1 hour at 37°C. 1 μL was plated onto small LB+ 100 μg/mL carbenicillin with 50% salt (LB Lennox Agar) for estimation of colony yields, and the remaining 1 mL was plated onto a single large LB+Carb with 50% salt.. Cells were washed the following day with 7 mL LB+Carb+25% glycerol and 1 mL of the wash used for plasmid extraction with the QIAprep Spin Miniprep Kit (Qiagen).

#### Step 2 plasmid library cloning

4 μg of the step 1 cloned library were cut with 4 μL BspQI (NEB) in a 200 μL reaction with 1X rCutSmart buffer at 50°C for 1 hour. Then, 4 μL calf-intestinal alkaline phosphatase (CIP, NEB) was added and the reaction resumed for 37°C for 1 hour. The expected 6294 bp band (only fragment) was gel-extracted and eluted in 50 μL EB, yielding 51 ng/μL of the cut step 1 library. The MAGESTIC 3.0 step 2 cloning insert (pF833) is 4754 bp and was amplified from pS1461 with KR1738 (5’-TCGATCGCTCTTCTgtttaagagctatgctggaaac-3’) and JS1530 (5’-GAATCTGAGTTACTGTCTGTTTTCCT-3’) in a 200 μL KAPA reaction (split into 4 x 50 μL aliquots in PCR tubes) with 10 ng pS1461 with the following program: 95°C 3 min, 30 cycles of [98°C 20 sec, 60°C 15 sec, 72°C 3 min], 72°C 5 min. pF833 was extracted from a 1% gel and eluted with 50 μL EB on two QIAquick columns each to yield 100 μL at 270 ng/μL. 4 μg of the pF833 elution were cut with 4 μL BspQI (NEB) in a 200 μL reaction with 1X rCutSmart buffer at 50°C for 1 hour, gel extracted from a 1% gel, and eluted with 50 μL EB on two QIAquick columns each to yield 100 μL at 22.1 ng/μL of the cut step 2 cloning insert (termed cV408). 50 ng of cV408 (4.6 kb) were ligated with 800 ng of the step 1 cut library (6.2 kb) with 1 μL T4 DNA ligase (NEB) in a 20 μL reaction in a thermocycler with the following program: 25°C 1 hour, 65°C 10 min, 4°C hold (heated lid at 50°C). The ligation was ethanol-precipitated, electroporated as described in the step 1 cloning section, and plated onto low-salt LB plates supplemented with 100 μg/mL carbenicillin and 30 μg/mL kanamycin (Kan). The plates were washed with 7 mL LB+Carb+Kan+25% glycerol with 1 mL of the wash used for plasmid extraction with the QIAprep Spin Miniprep Kit (Qiagen) to yield the step 2 plasmid library.

#### MAGESTIC 3.0 multiplexed editing in yeast

First, 4 μg of the step 2 plasmid library was cut with 4 μL I-SceI in 1X rCutSmart buffer in 200 μL total volume at 37°C for 1 hour, extracted from a 1% gel, and eluted in 100 μL EB on a single QIAquick column to yield 24.50 ng/μL of the linearized step 2 guide-donor library. This removes the intermediate placeholder p*GalL-SceI-kanR*-*HIS3* insert from the step 2 library and provides homology overlaps for *in vivo* plasmid assembly with a barcoded insert (described below), including a 260 bp overlap of the *CYC1* terminator region adjacent to the sgRNA and a 77 bp overlap consisting of the synthetic SaCas9 target site (X) and Ec86 retron msr/msd sequences flanking the donor DNA.

The MAGESTIC 3.0 barcoded insert (pF835) was prepared in two steps from plasmid pS1460. First, it was amplified by KR2132 forward primer (500 nM, 5’-CTcctgcaggCAAATTAAAGC-3’) and KR2133 reverse primer (500 nM, 5’-GGAtaccgTTCGTATAGCATACATTATACGAAGTTATgcacgTatccTAGGGNNNNNNNNNNNNNNNNNNNNGAGCATGACCTGTCGACG-3’) in a 200 μL KAPA HiFi HotStart reaction with 50 ng pS1460 template with the following program: 95°C 3 min, 30 cycles of [98°C 20 sec, 60°C 15 sec, 72°C 4 min], 72°C 5 min. The resulting product (pF834) was extracted from a 1% gel as a 4413 bp fragment and eluted on 2 separate columns in 50 μL EB each to yield 138.80 ng/μL. 1 μg of pF834 (∼7 μL) was used a template in an 800 μL KAPA reaction with the KR2132 forward primer (500 nM, 5’-CTcctgcaggCAAATTAAAGC-3’), an inner reverse primer KR2134 (present at 50 nM; 5’-TGCGCACCTGAGTGGAAGAAACCCACGGAGGAGTGGGTAaggATGGATAGTGCCCTATGAAATAGGGGAtaccgTTCGTATAGCATACAT-3’) and an outer reverse primer KR2135 (500 nM; 5’-gctcttcaaacAGGAAACCCGTTTCTTCTGACGTAAGGGTGCGCACCTGAGTGGAAGAAACCCACGGAGGAGT-3’) with the following program: 95°C 3 min, 30 cycles of [98°C 20 sec, 60°C 15 sec, 72°C 4 min], 72°C 5 min. The resulting product (pF835) was extracted from a 1% gel as a 4518 bp fragment and eluted on 4 separate columns in 100 μL EB each to yield 400 μL at 20 ng/μL.

The pF835 insert contains the 260 bp *CYC1* terminator region for homology to the linearized step 2 guide-donor library (described above), a galactose-inducible I-SceI, which cuts an I-SceI site present on the guide-donor plasmid as well as two sites flanking the editing machinery integrated at the REDI locus, a *kanR* marker which provides for ∼880 bp homology during integration of the barcode into the barcode locus post-editing, a *HIS3* marker for selection after yeast transformation, a *GAL7* terminator for the retron donor ncRNA, a 20-mer random barcode (bc1) for tracking edits, an ∼80 bp region consisting of the lox66 variant site and surrounding region which serves as homology for barcode locus integration after editing, and finally the 77 bp region serving as homology to the linearized step 2 guide-donor library.

For the multiplexed editing, 8 mL of yKR1026 was cultured in 2X YPAD from OD 0.1 to OD 0.8. The culture was spun down at 2500 g for 5 minutes and washed with 1 mL H2O. 320 μL 50% PEG, 48 μL 1M LiOAc, 40 μL 10 mg/mL ssDNA were added to the cell pellet along with 1 μg of the linearized step 2 guide-donor library (52 μL) and 1 μg of pF835 (50 μL). The transformation mix was vortexed vigorously to fully resuspend the cell pellet and then split into 4 replicate PCR tubes. The tubes were incubated at 42°C for 1 hour, spun down at 5000 g for 1 minute, resuspended in 100 μL 0.5 M sorbitol and then plated onto SC-Ura-His plates and incubated at 30°C for 3 days. Cells were washed from the plates in 3 mL SC-Ura-His +15% glycerol, combined, and then split into two replicate yeast libraries (denoted yL435 and yL436). These libraries were inoculated into 700 μL liquid SC-Ura-His and outgrown in a custom yeast grower system developed by Tecan US which consists of 4 Infinite 200 Pro M Nano microplate readers (Tecan Austria GmbH; firmware downgraded to version 5.26 to allow for vigorous shaking with the Pegasus yeast grower system), four 4°C RIC20 remote-controlled chilling/heating block (Torrey Pines Scientific) and a EVO 150 liquid handler (Tecan). The cultures were started at OD 0.1 in 48-well plates (∼equivalent to OD 0.3 in a 1 cm cuvette) and shaken at 30°C with constant linear shaking (6 mm amplitude) with OD readings performed at intervals of 5 min. Upon reaching a trigger OD of 0.76, the system automatically transferred 11 μL of culture into 689 μL SC-His + galactose + 1 μg/mL aTc to induce the barcode integration and guide-donor plasmid removal (1/64 of the 700 μL culture volume transferred to allow for ∼6 generations). Importantly, the yeast grower system also saves the remaining 600 μL of the culture by transferring onto separate 48-well plate situated on the 4°C chilled blocks. This was repeated for a 2^nd^ barcoding outgrowth, followed by a final outgrowth in SC-His + dextrose + 300 μM 5-fluorocytosine (5FC) + 1 mg/mL 5-fluoroorotic acid (5FOA) for counter-selection to ensure residual guide-donor plasmid by virtue of the *FCY1* marker present on the pS1439 backbone and to ensure completion of barcoding by virtue of the *URA3* marker present with the editing machinery cassette at the barcode locus. For one additional outgrowth, 44 μL of the 5FC/5FOA outgrowths were transferred to 4 mL YPD+G418 + 1 mM 5FC and grown overnight. The following day, 3 mL of this outgrowth was combined with 3 mL 50% glycerol to yield six 1 mL aliquots of edited and barcoded libraries. The OD600 of these glycerol stocks were ∼4.0. To prepare for a replicate of the screen, a 1 mL aliquot was inoculated in 10 mL YPD+G418 for overnight outgrowth.

### Fitness analysis

To assess barcode abundance over IAA treatment (i.e., Nrd1 depletion), we diluted overnight cultures of the library (yL434 for the REDI-parsed NNS library, or yL435/yL436 for the repeat Nrd1 library) to ∼OD600 0.5, in 700 μL YPD in 48-well plates. The first row contains YPD, and each subsequent well of each column contained YPD, or YPD supplemented with 0.5 uM IAA, or condition specific media supplemented with 0.5 uM IAA. Plates were incubated in the yeast grower system described above, except transferring 1/16th (44 μL out of 700 μL) of the content of the culture to the subsequent well to allow for 4 generations of growth for each passage for a total of 5 passages, or 20 generations of growth. 600 μL of the remaining culture was transferred to a separate plate on a 4°C RIC20 remote-controlled chilling/heating block (Torrey Pines Scientific) until the end of the experiment.

gDNA was purified using a Masterpure Yeast DNA Purification (Epicentre) following the manufacturer’s instructions. PCRs with primers containing indexing sequences were then performed in 20 μL reactions to amplify the genomic-integrated barcodes. The libraries were sequenced on the AVITI system (Element Biosciences; sequencing conducted by the Stanford Human Immune Monitoring Center genomics core facility) with 2x150 bp reads and reads processed with custom scripts (see workflow and scripts at https://github.com/k-roy/MAGESTIC/tree/master/NRD1). gDNA was also extracted from cells from the plate wash and editing outgrowth steps (from SC-Ura-His) to allow for sequencing of the bc1-donor-bc0 region from the yeast plasmid assembly to enable associating each bc1 with a guide-donor cassette.

Normalized barcode counts were analyzed across generations, treatments (YPD, YPD no IAA), and replicates. Edits were annotated as missense, synonymous, PTC, or deletion. Counts were stabilized by adding a pseudocount and, for each barcode, scaled to its mean at generation 0; the fitness index was defined as log2 fold-change relative to generation 0. For each amino-acid position × mutation type × treatment, barcode trajectories were collapsed to a mean fitness-over-time series. To avoid late low-signal noise, we trimmed each collapsed series at the first occurrence of sustained depletion (two consecutive post-G0 points below a fold-change cutoff), and estimated a linear slope of log2 fold-change versus generation within this window. Slopes were median-centered within each treatment to obtain adjusted slopes. Hit calling compared YPD to YPD noIAA using three criteria: (i) the YPD adjusted slope lay in the negative tail of the YPD distribution (5th percentile), (ii) the YPD effect was distinctly stronger than YPD_noIAA (≥3x more negative), and (iii) the variant showed endpoint depletion in YPD (final fold-change <0.1). Barcode-level slopes, computed within the same trimmed window, were used to summarize support (number and fraction of barcodes independently meeting the criteria).

### Microscopy

Cells were grown in exponential phase (0.5 ≤ OD600 ≤ 1) in CSM medium at 30°C and treated with IAA at a final concentration of 500 μM for 1 h before imaging. Wide-field fluorescence images were acquired using a Leica DM6000B microscope with a 100X/1.4 NA (HCX Plan-Apo) oil immersion objective and a CCD camera (CoolSNAP HQ; Photometrics). The acquisition system was piloted by the MetaMorph software (Molecular Devices). Images were scaled equivalently and merged using ImageJ. An average of three experiments, each of them visualizing at least 300 cells per condition, is shown in the figures.

### RNA extraction

Cells were grown in logarithmic phase, and 6 OD_600_ worth of cells were collected. Total RNAs were extracted by resuspending cell pellets in 1 volume of acidic Phenol (pH 4.3, Sigma) supplemented with 1 volume of AES Buffer (50 mM Sodium Acetate pH 5.5, 10 mM EDTA, 1% SDS). Mixtures were incubated at 70°C with shaking (2000 RPM) for 30 minutes in a Thermomixer (Eppendorf), before being spanned at 20000 g, 4°C for 10 minutes. Aqueous phases were recovered and subjected to one extra round of hot acidic Phenol extraction followed by one round of Chloroform extraction. Total RNAs were finally precipitated with absolute Ethanol and Sodium Acetate pH 5.5, washed once with 70% Ethanol, dried and resuspended in 30μL of RNAse-free H2O. 60 to 120μg of total RNAs were recovered in routine

### Reverse transcription and quantitative PCR

Reverse transcription was performed using random hexamers primers annealing at multiple *loci* in the *S. cerevisiae* genome. 4 μg of total RNAs were mixed to 200 ng of random hexamers in a 20 μL reaction containing 50 mM Tris-HCl pH 8.3, 75 mM KCl, 3 mM MgCl2, and 5 mM DTT. Samples were first incubated for 15 min at 70°C to allow RNA denaturation. Then temperature was slowly decreased to 37°C to allow annealing of primers. Lastly, synthesis of cDNAs was performed by adding 200 Units of MLV-Reverse Transcriptase (Invitrogen) for 45 minutes at 37°C.

To assess the amount of cDNAs reverse transcribed, quantitative PCR (qPCR) was carried out using different primer pairs for each target (SNR13, NEL250c, ACT1). Specifically, qPCR was performed in a 20 μL reaction using LUNA qPCR Master Mix (New England Biolabs) following manufacturer instruction.

### Total protein lysates and Western Blotting analysis

Cells were grown in logarithmic phase, and 6 OD_600_ worth of cells were pelleted. Total cell extract was obtained by lysing each cell pellet in NaOH 0.25 M, 1% 2-Mercaptoethanol and precipitating proteins in TCA 6%. Protein pellets were then resuspended in LDS Sample Buffer (Invitrogen) and incubated at 95°C for 5 minutes with shaking.

Samples were loaded onto a 4-12 % SDS-PAGE gels (Thermo) and migrated for 1 hour at 120V then transferred onto a nitrocellulose membrane at 130V for 1.5 hours at 4°C. Membrane was first stained with Ponceau S and scanned, then washed in TBSt 1X (Tris-Borate-Sodium-Tween). Subsequently, membrane was blocked in Buffer TBSt 1X supplemented with 5% of dry non-fat milk, then incubated overnight with primary antibodies (α-GFP, NeuroMab) in Buffer TBSt 1X, 5% dry non-fat milk. Next, excess of primary antibody was first washed away with TBSt 1X, then membrane was incubated for 1 hour at 22°C with a secondary antibody coupled to HRP (α-mouse HRP conjugate). Finally, excess of secondary antibody was washed away with TBSt 1X, and membranes were revealed using SuperSignal West Dura Chemoluminescent Substrate (Thermo) on Chemidoc scanner (Biorad).

## Data and code availability

Code for processing short-read NGS amplicons (guide-donor-bc0, bc1-donor-bc0, and genome-integrated bc1) are available at https://github.com/k-roy/MAGESTIC/tree/master/NRD1. All NGS data produced by this study will be made available upon publication.

## ACKNOWLEDGMENTS

U.A. was supported by a Stanford University School of Medicine Dean’s fellowship and by an EMBO long-term postdoctoral fellowship (ALTF 889-2022).

---

**Supplementary Figure 1 related to Figure 1.**
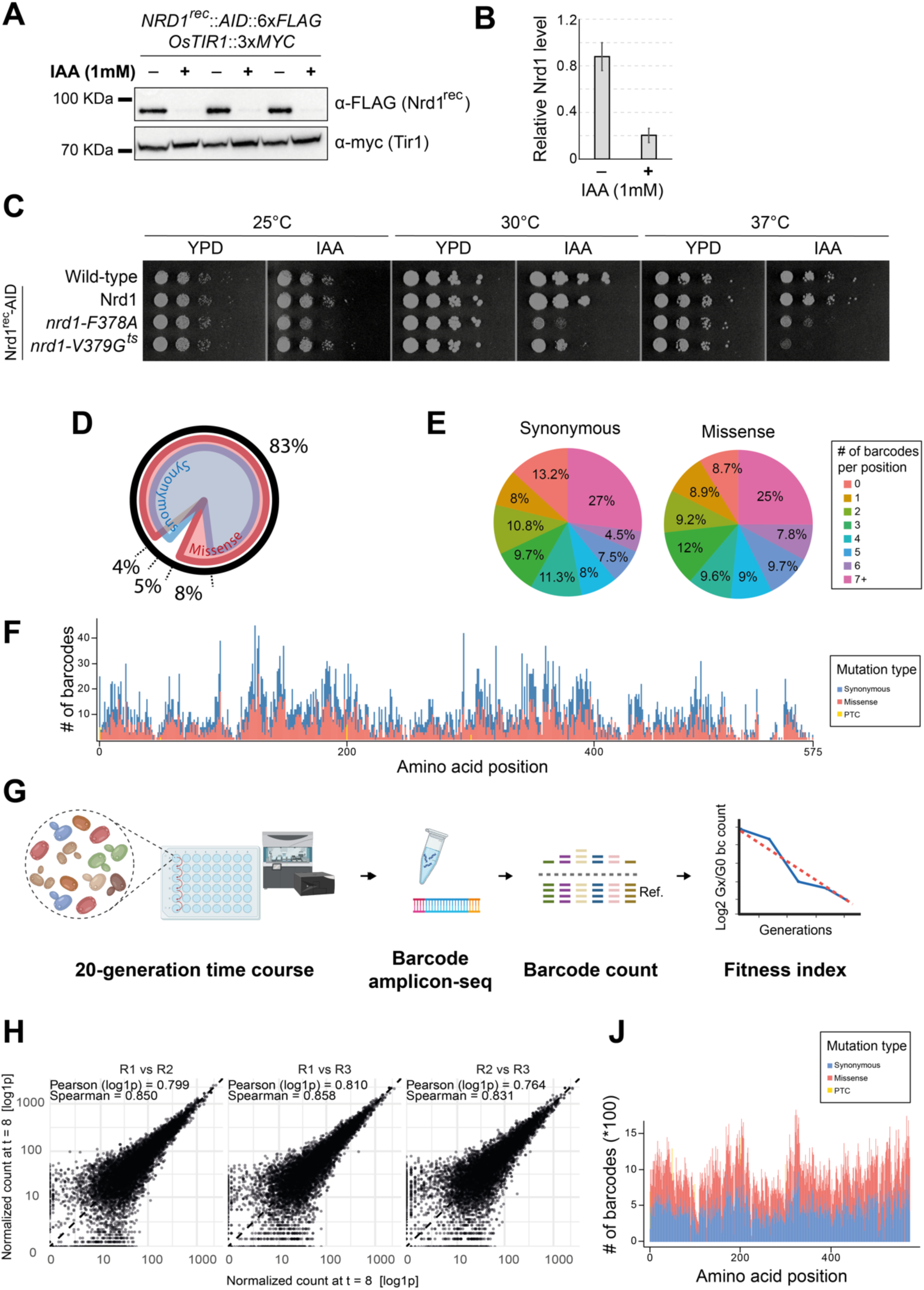
**A.** Western blotting analysis of the Nrd1rec expression level in presence and absence of IAA treatment (1 hour). The detected levels of Tir1 are also shown. **B.** Quantification of A and other 2 independent replicates. **C.** Growth test of plates of the indicated strains in presence and absence of IAA and at the indicated temperature. Plates were incubated for 2 days before being photographed. **D.** Pie plot representation of the percentages and the overlap of the edits present in the Nrd1 alanine scanning library compared to the total amount of residues in Nrd1. **E.** Pies of the percentages of amino acid positions in Nrd1 associated to the indicated numbers of barcodes. **F.** Distribution of the edits along the Nrd1 gene. Mutation categories (missense, synonymous and premature termination codons, PTCs) are shown with different colors as indicated in the legend. **G.** Schematic representation of the 20-generation time course experiment and the downstream fitness score quantification. Cells are grown for 20 generations in an automatized platform, with samples collected each four generations (see more in Materials and Methods). After gDNA extraction and PCR amplification of the barcode *locus*, the products are sequenced and the normalized count of barcodes quantified along the time course to fit a linear regression, the slope of which is used as a fitness index. **H.** Dispersion plots of the of the barcode counts for the indicated replicates. Correlation scores are shown for each replicate pair. **J.** Edit distribution along the *NRD1* gene from barcode counts during library prep (pre-editing stage).

**Supplementary Figure 2 related to Figure 1.**
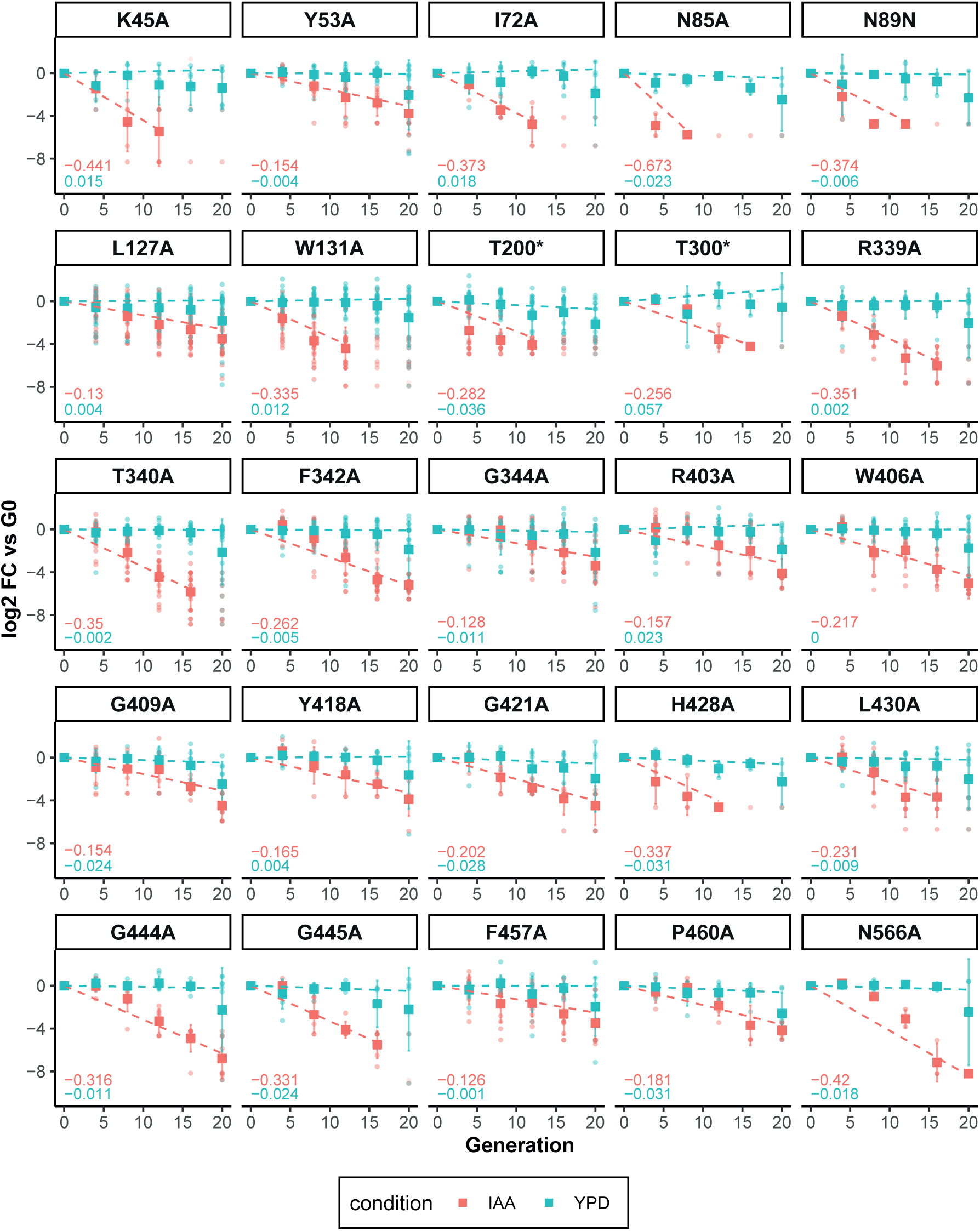
Log2 FC (Gn vs G0) of the normalized barcode counts of the indicated edits. The linear regression is shown as a dashed line. The computed slopes are shown at the bottom-left of each plot following the indicated color-code.

**Supplementary Figure 3 related to Figure 2.**
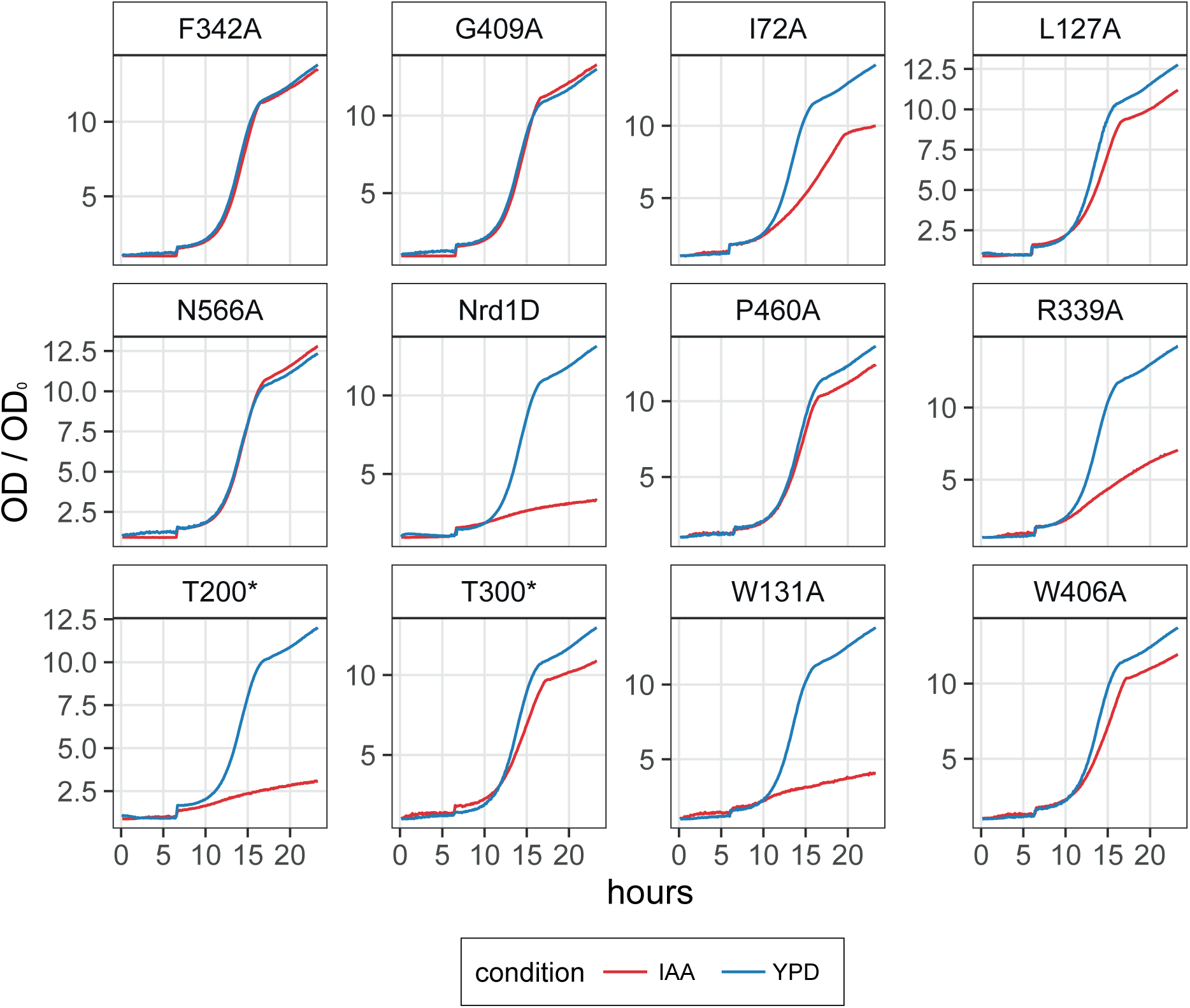
Division rate (OD_t_/OD_0_) of the indicated strains in non-competitive growth assays, with and without IAA treatment.

**Supplementary Figure 4 related to Figure 3.**
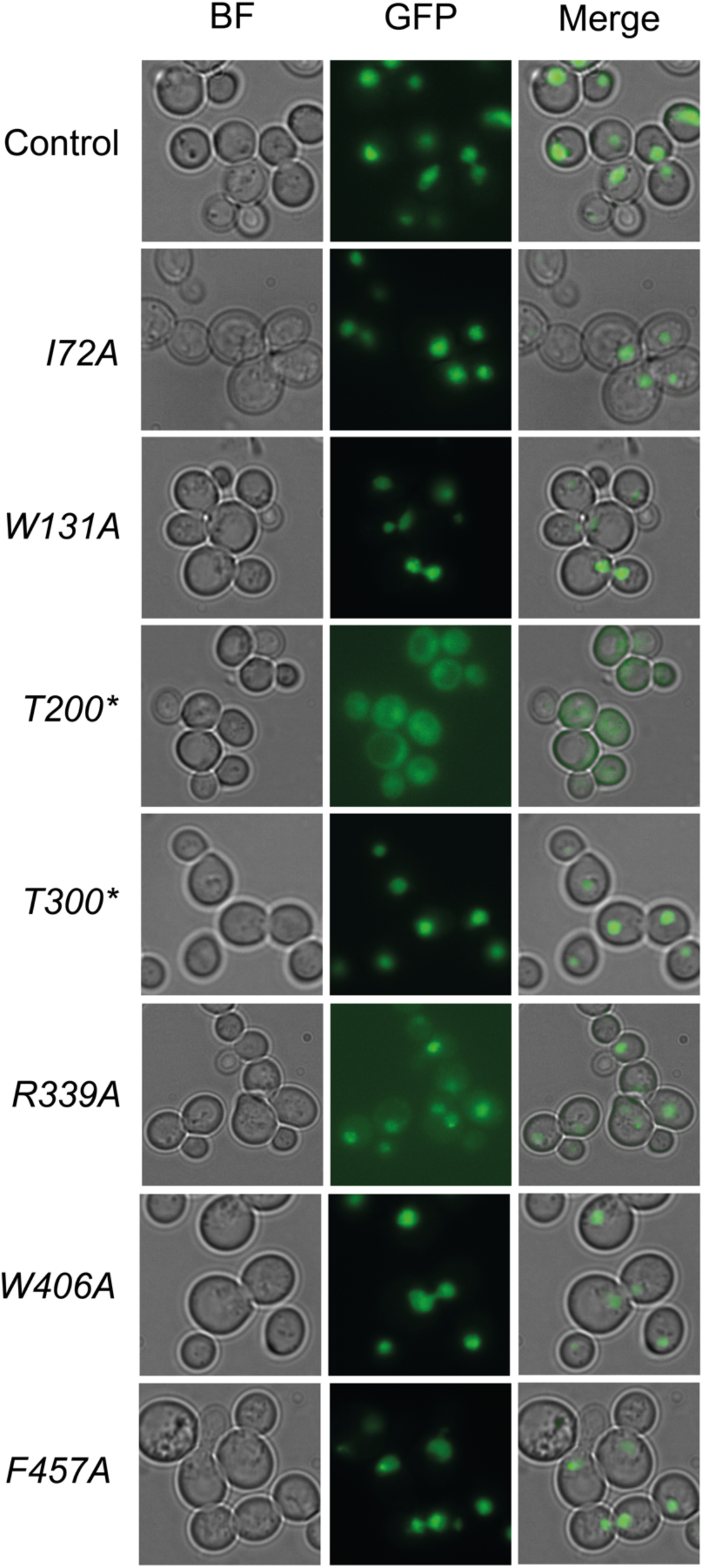
Representative microscopy fields of the indicated strains.

## Notes

https://github.com/k-roy/MAGESTIC/tree/master/NRD1

### Competing Interest Statement

The authors have declared no competing interest.

## REFERENCES

Aiello U, Challal D, Wentzinger G, Lengronne A, Appanah R, et al. 2022. Sen1 is a key regulator of transcription-driven conflicts. Mol. Cell. 82(16):2952–2966.e6

Araya CL, Fowler DM. 2011. Deep mutational scanning: assessing protein function on a massive scale. Trends Biotechnol. 29(9):435–42

Arigo JT, Eyler DE, Carroll KL, Corden JL. 2006. Termination of Cryptic Unstable Transcripts Is Directed by Yeast RNA-Binding Proteins Nrd1 and Nab3. Mol. Cell. 23(6):841–51

Azizoglu A, Brent R, Rudolf F. 2021. A precisely adjustable, variation-suppressed eukaryotic transcriptional controller to enable genetic discovery. eLife. 10:e69549

Bacikova V, Pasulka J, Kubicek K, Stefl R. 2014. Structure and semi-sequence-specific RNA binding of Nrd1. Nucleic Acids Res. 42(12):8024–38

Conrad NK, Wilson SM, Steinmetz EJ, Patturajan M, Brow DA, et al. 2000. A yeast heterogeneous nuclear ribonucleoprotein complex associated with RNA polymerase II. Genetics. 154(2):557–71

Cooper S, Obolenski S, Waters AJ, Bassett AR, Coelho MA. 2024. Analyzing the functional effects of DNA variants with gene editing. Cell Rep. Methods. 4(5):

Cunningham BC, Wells JA. 1989. High-Resolution Epitope Mapping of hGH-Receptor Interactions by Alanine-Scanning Mutagenesis. Science. 244(4908):1081–85

Darby MM, Serebreni L, Pan X, Boeke JD, Corden JL. 2012. The Saccharomyces cerevisiae Nrd1-Nab3 Transcription Termination Pathway Acts in Opposition to Ras Signaling and Mediates Response to Nutrient Depletion. Mol. Cell. Biol. 32(10):1762–75

DiCarlo JE, Norville JE, Mali P, Rios X, Aach J, Church GM. 2013. Genome engineering in Saccharomyces cerevisiae using CRISPR-Cas systems. Nucleic Acids Res. 41(7):4336–43

Findlay GM, Daza RM, Martin B, Zhang MD, Leith AP, et al. 2018. Accurate classification of BRCA1 variants with saturation genome editing. Nature. 562(7726):217–22

Flagfeldt DB, Siewers V, Huang L, Nielsen J. 2009. Characterization of chromosomal integration sites for heterologous gene expression in Saccharomyces cerevisiae. Yeast Chichester Engl. 26(10):545–51

Franco-Echevarría E, González-Polo N, Zorrilla S, Martínez-Lumbreras S, Santiveri CM, et al. 2017. The structure of transcription termination factor Nrd1 reveals an original mode for GUAA recognition. Nucleic Acids Res. 45(17):10293–305

Gould SI, Wuest AN, Dong K, Johnson GA, Hsu A, et al. 2023. High throughput evaluation of genetic variants with prime editing sensor libraries

Han Z, Jasnovidova O, Haidara N, Tudek A, Kubicek K, et al. 2020. Termination of non-coding transcription in yeast relies on both an RNA Pol II CTD interaction domain and a CTD-mimicking region in Sen1. EMBO J.

Han Z, Libri D, Porrua O. 2017. Biochemical characterization of the helicase Sen1 provides new insights into the mechanisms of non-coding transcription termination. Nucleic Acids Res. 45(3):1355–70

Harvey CJB, Tang M, Schlecht U, Horecka J, Fischer CR, et al. HEx: A heterologous expression platform for the discovery of fungal natural products. Sci. Adv.

Heo D, Yoo I, Kong J, Lidschreiber M, Mayer A, et al. 2013. The RNA polymerase II C-terminal domain-interacting domain of yeast Nrd1 contributes to the choice of termination pathway and couples to RNA processing by the nuclear exosome. J. Biol. Chem. 288(51):36676–90

Howard MK, Hoppe N, Huang X-P, Mitrovic D, Billesbølle CB, et al. 2025. Molecular basis of proton sensing by G protein-coupled receptors. Cell. 188(3):671–687.e20

Inoue F, Kircher M, Martin B, Cooper GM, Witten DM, et al. 2017. A systematic comparison reveals substantial differences in chromosomal versus episomal encoding of enhancer activity. Genome Res. 27(1):38–52

Jasin M, Rothstein R. 2013. Repair of Strand Breaks by Homologous Recombination. Cold Spring Harb. Perspect. Biol. 5(11):a012740–a012740

Jensen TH, Jacquier A, Libri D. 2013. Dealing with Pervasive Transcription. Mol. Cell. 52(4):473–84

Kubicek K, Cerna H, Holub P, Pasulka J, Hrossova D, et al. 2012. Serine phosphorylation and proline isomerization in RNAP II CTD control recruitment of Nrd1. Genes Dev. 26(17):1891–96

Langdon EM, Qiu Y, Ghanbari Niaki A, McLaughlin GA, Weidmann CA, et al. 2018. mRNA structure determines specificity of a polyQ-driven phase separation. Science. 360(6391):922–27

Maharana S, Wang J, Papadopoulos DK, Richter D, Pozniakovsky A, et al. 2018. RNA buffers the phase separation behavior of prion-like RNA binding proteins. Science. 360(6391):918–21

Montserrat-Canals M, Cordara G, Krengel U. 2025. Allostery. Q. Rev. Biophys. 58:e5

Morawska M, Ulrich HD. 2013. An expanded tool kit for the auxin-inducible degron system in budding yeast. Yeast. 30(9):341–51

Nishimura K, Fukagawa T, Takisawa H, Kakimoto T, Kanemaki M. 2009. An auxin-based degron system for the rapid depletion of proteins in nonplant cells. Nat. Methods. 6(12):917–22

O’Rourke TW, Loya TJ, Head PE, Horton JR, Reines D. 2015. Amyloid-like assembly of the low complexity domain of yeast Nab3. Prion. 9(1):34–47

Papagiannakis A, Niebel B, Wit EC, Heinemann M. 2017. Autonomous Metabolic Oscillations Robustly Gate the Early and Late Cell Cycle. Mol. Cell. 65(2):285–95

Porrua O, Libri D. 2013. A bacterial-like mechanism for transcription termination by the Sen1p helicase in budding yeast. Nat. Struct. Mol. Biol. 20(7):884–91

Porrua O, Libri D. 2015. Transcription termination and the control of the transcriptome: why, where and how to stop. Nat. Rev. Mol. Cell Biol. 16(3):190–202

Roy KR, Smith JD, Li S, Vonesch SC, Nguyen M, et al. 2024. Dissecting quantitative trait nucleotides by saturation genome editing

Roy KR, Smith JD, Vonesch SC, Lin G, Tu CS, et al. 2018. Multiplexed precision genome editing with trackable genomic barcodes in yeast. Nat. Biotechnol. 36(6):512–20

Schaughency P, Merran J, Corden JL. 2014. Genome-Wide Mapping of Yeast RNA Polymerase II Termination. PLoS Genet. 10(10):e1004632

Schmid CM, Gregor A, Ruiz A, Manso Bazús C, Herman I, et al. 2025. Further delineation of the SCAF4-associated neurodevelopmental disorder. Eur. J. Hum. Genet. 33(5):588– 94

Schulz D, Schwalb B, Kiesel A, Baejen C, Torkler P, et al. 2013. Transcriptome Surveillance by Selective Termination of Noncoding RNA Synthesis. Cell. 155(5):1075–87

Schymkowitz J, Borg J, Stricher F, Nys R, Rousseau F, Serrano L. 2005. The FoldX web server: an online force field. Nucleic Acids Res. 33(suppl_2):W382–88

Smith JD, Schlecht U, Xu W, Suresh S, Horecka J, et al. 2017. A method for high-throughput production of sequence-verified DNA libraries and strain collections. Mol. Syst. Biol. 13(2):

Starr TN, Greaney AJ, Hilton SK, Ellis D, Crawford KHD, et al. 2020. Deep Mutational Scanning of SARS-CoV-2 Receptor Binding Domain Reveals Constraints on Folding and ACE2 Binding. Cell. 182(5):1295–1310.e20

Steinmetz EJ, Brow DA. 1998. Control of pre-mRNA accumulation by the essential yeast protein Nrd1 requires high-affinity transcript binding and a domain implicated in RNA polymerase II association. Proc. Natl. Acad. Sci. 95(12):6699–6704

Thiebaut M, Kisseleva-Romanova E, Rougemaille M, Boulay J, Libri D. 2006. Transcription Termination and Nuclear Degradation of Cryptic Unstable Transcripts: A Role for the Nrd1-Nab3 Pathway in Genome Surveillance. Mol. Cell. 23(6):853–64

Tudek A, Porrua O, Kabzinski T, Lidschreiber M, Kubicek K, et al. 2014. Molecular Basis for Coordinating Transcription Termination with Noncoding RNA Degradation. Mol. Cell. 55(3):467–81

Vasiljeva L, Kim M, Mutschler H, Buratowski S, Meinhart A. 2008. The Nrd1–Nab3–Sen1 termination complex interacts with the Ser5-phosphorylated RNA polymerase II C-terminal domain. Nat. Struct. Mol. Biol. 15(8):795–804

Yuryev A, Patturajan M, Litingtung Y, Joshi RV, Gentile C, et al. 1996. The C-terminal domain of the largest subunit of RNA polymerase II interacts with a novel set of serine/arginine-rich proteins. Proc. Natl. Acad. Sci. 93(14):6975–80

Zhang Y, Chun Y, Buratowski S, Tong L. 2019. Identification of Three Sequence Motifs in the Transcription Termination Factor Sen1 that Mediate Direct Interactions with Nrd1. Structure. 27(7):1156–1161.e4

